# scSketch: Interactive Sketch-based Trajectory Exploration and Pathway-Aware Analysis of Single-Cell Data

**DOI:** 10.64898/2026.04.16.718997

**Authors:** Askar Temirbek, Fritz Lekschas, Kris Sankaran, Andres Colubri

## Abstract

Interactively exploring gene expression gradients across low-dimensional cell embeddings is central to single-cell RNA sequencing analysis, yet there aren’t tools that allow users to sketch trajectories and interactively compute pathway-level interpretation. We present scSketch, a tool that enables users to iteratively explore and test trajectory hypotheses in single-cell data while maintaining statistical validity and biological interpretability. Specifically, users apply *interactive directional sketching* to draw trajectories across embeddings and probe continuous processes such as cellular differentiation and cell state transitions. scSketch automatically computes gene-trajectory correlations and applies online false discovery rate (FDR) control to maintain statistical validity during iterative exploration. Significant genes are grouped into Reactome pathways for contextual interpretation. Applied to human oral keratinocytes infected with human cytomegalovirus, scSketch revealed infection-associated gradients involving interferon responses, metabolic remodeling and autophagy. Together, these features position scSketch as a bridge between exploratory visualization and mechanistic insight in single-cell biology.

## 1. Introduction

### Single-cell RNA-sequencing analysis and its goals

Single-cell RNA sequencing (scRNA-seq) measures gene expression at individual cell resolution, enabling the characterization of cellular heterogeneity within complex tissues and systems [1]. Key goals in analyzing scRNA-seq data include identifying distinct cell types and states, as well as resolving continuous transitions between them, such as developmental trajectories, differentiation programs, and gradual response gradients [2].

### Standard single-cell workflows

In common practice, scRNA-seq analyses follow a programmatic pipeline involving quality control, dimensionality reduction, graph-based clustering, and visualization in low-dimensional embeddings [3-5]. Downstream interpretation typically relies on differential expression (DE) and pathway enrichment applied post hoc to cluster-defined gene lists. Popular environments such as Seurat [3] and Scanpy [4] streamline these workflows and provide basic interactivity (e.g., selecting cells, recoloring plots), but generally decouple exploratory selection from immediate pathway context and do not natively support iterative, user-driven hypothesis testing with sequential error control. scSketch in context. scSketch addresses these limitations by enabling interactive exploration of continuous structure directly within low-dimensional embeddings. Users visually define candidate trajectories or regions of interest and immediately quantify how gene expression varies along these user-specified directions, while maintaining statistical validity through online false discovery rate (FDR) control. Results are presented inline with pathway-level interpretation via Reactome, allowing exploratory gestures to yield mechanistic insight without leaving the notebook. In contrast to algorithm-defined trajectory inference methods that produce a fixed ordering, scSketch supports flexible, hypothesis-driven interrogation of visually apparent gradients and alternative directions of variation.

### Contributions

We present scSketch, an open-source interactive visualization and analysis software tool for the scverse ecosystem. Improving upon a previous prototype developed by one of the authors [6], scSketch introduces three key innovations:

i. Free-hand interactive directional sketching (IDS) - users can draw linear or non-linear vectors (as a brush to cover user-specified areas of target cells) directly in the embedding to define putative biological trajectories.
ii. Online FDR control - statistical rigor is maintained during iterative sketching by applying online FDR methods to gene-trajectory associations.
iii. Pathway-aware interpretation - integration with *Reactome* [7] allows users to group correlated genes into shared pathways and view molecular diagrams, bridging exploratory selection with biological context.

Built on the GPU-accelerated *Jupyter Scatter* [8] and the *anywidget* [9] framework, scSketch supports high-performance rendering [10] and seamless Jupyter integration, allowing single-cell datasets as large as 20 million cells to be explored interactively without leaving the notebook environment. We demonstrate its utility on scRNA-seq data from human oral keratinocytes (HOKs) infected with human cytomegalovirus (HCMV), reproducing known infection-associated gradients and revealing additional host-response pathways [11]. These results illustrate how scSketch transforms exploratory sketches into statistically robust and biologically meaningful insights, complementing traditional DE and pseudotime analyses. scSketch is open source and available as a Python package via PyPI (https://pypi.org/project/scsketch/), with source code and documentation hosted on GitHub (https://github.com/colabobio/scsketch).

## 2. Methods

### 2.1. Software architecture and integration

scSketch is implemented as an interactive Python package that runs inside Jupyter notebooks. The system is built as a composition of anywidget components, with Jupyter Scatter handling GPU-accelerated rendering of low-dimensional embeddings and capture of user interactions, including cell selection and free-hand sketching. Other custom anywidget user interface components, including ranked gene lists and Reactome pathway displays, are combined into the scSketch object that users can access from their notebooks. These components manage gene lists, statistical summaries, and pathway visualization, enabling tight integration between interactive exploration and downstream analysis. **Figure 1A** shows scSketch’s architecture across server side (Jupyter server) and front end (web browser) modules.

**Figure 1.**
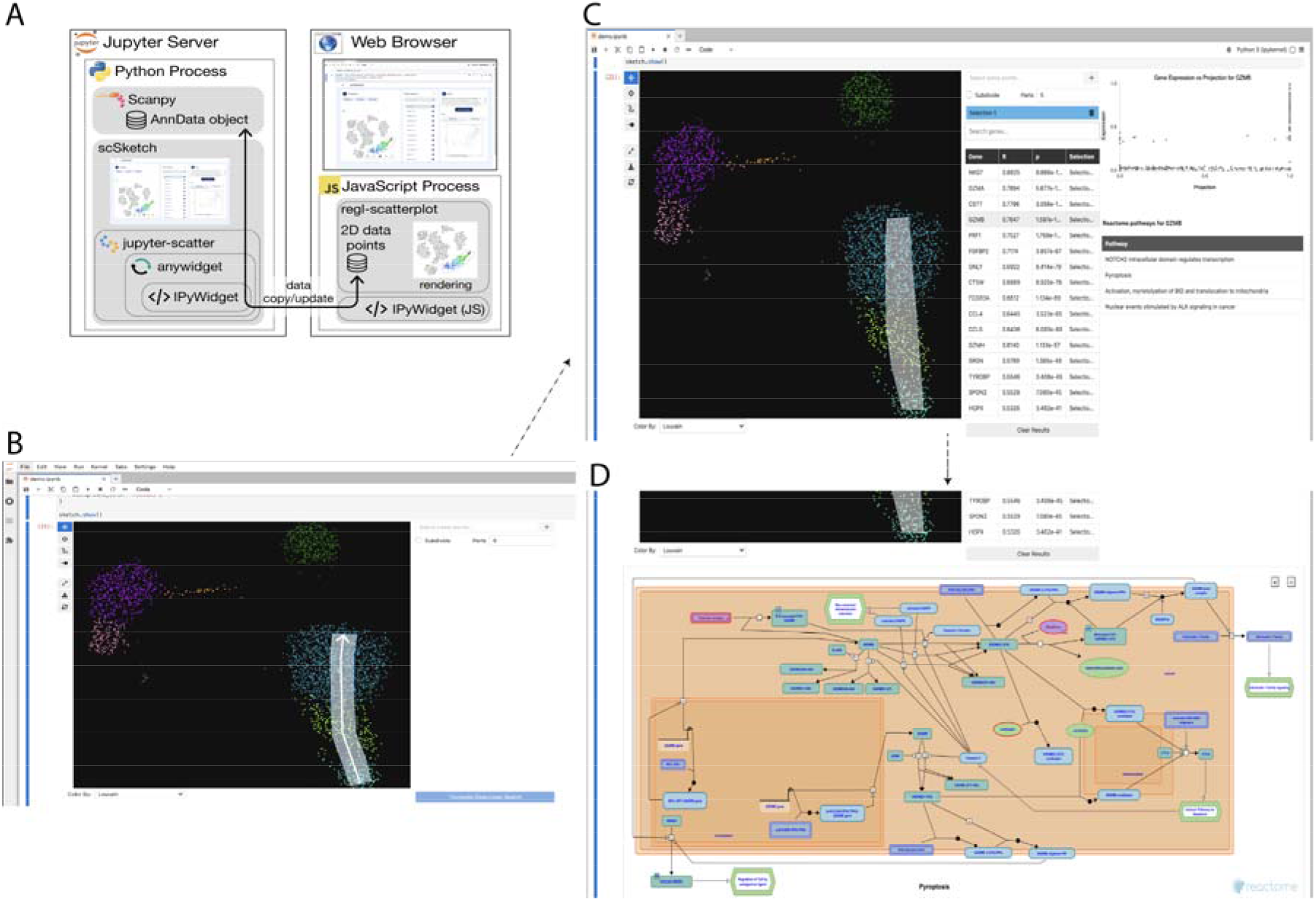
Architecture and interactive workflow of scSketch. **(A)** Overview of scSketch architecture. The tool runs as a Python object within the Jupyter server for data access and computations, while interactive visualization is handled by the GPU-accelerated regl-scatterplot library in the browser. **(B)** Jupyter notebook environment showing scSketch applied to a scRNA-seq PBMC dataset from Hao et al. (2021), visualized using a UMAP embedding with aggregated cell-type labels. scSketch is operating in brush mode, drawing a desired direction with the target cells inside the brush selection (brush selection and direction of drawing highlighted in white color for clarity). **(C)** Saving the selected direction with target cells with a button click, followed by clicking the Calculate Directional Search button to perform directional analysis. This produces a ranked gene list with Pearson correlations, p-values, and selection associated with the results. Clicking a gene from the directional results displays a scatter plot of its expression alongside the drawn direction and the associated Reactome pathways in a table. **(D)** Upon pressing a desired Reactome pathway, a pathway diagram appears that shows the pathway within which the target gene product belongs. Alt text: Four-panel figure showing scSketch’s architecture and interactive workflow in a Jupyter notebook. **(A)** Diagram of the system split between a Python process on the Jupyter server (handling data access and computation with AnnData/Scanpy and scSketch widgets) and a web browser front end (rendering the embedding with a GPU-accelerated scatterplot and sending user interactions back to Python). **(B)** Screenshot of a PBMC UMAP in scSketch with a white brush stroke indicating a user-drawn directional selection. **(C)** Screenshot of the directional analysis results table listing genes with correlations and p-values, with a gene selected to view expression along the drawn direction and associated Reactome pathways. **(D)** Screenshot of a Reactome pathway diagram displayed after selecting a pathway from the table.

scSketch is deeply integrated with the scverse ecosystem. Users can load AnnData objects, visualize embeddings like UMAP [12], define trajectories or regions of interest by drawing on embeddings, and color cells by metadata or gene expression, all within their code notebook. Since scSketch is ultimately an IPyWidget, data derived from interactive exploration - such as lists of selected genes and associated statistics - are immediately accessible for downstream analyses. This design avoids the tool switching and context fragmentation common in current workflows that separate interactive visualization from statistical analysis [13, 14], enabling rapid hypothesis generation within a single workflow.

### 2.2. User workflow

The overall workflow of scSketch is illustrated in **Figure 1B-D**. Users initialize scSketch by passing an existing AnnData object, after which a two-dimensional embedding is rendered in an interactive scatterplot. To investigate continuous structure in the data, users apply IDS to draw directional sketches or regions of interest directly onto the embedding using interactive selection tools. Each sketch defines a user-specified direction or trajectory that is immediately converted into a quantitative representation linking cell positions to gene expression values. Sketch-derived results, including ranked gene lists and statistical summaries, remain available programmatically for further analysis without leaving the notebook. Multiple testing arising from repeated directional sketching is handled using an online FDR framework [15,16]. In addition to directional sketching, scSketch provides a DE mode for discrete comparisons: users draw a free-form lasso to select an arbitrary cell set and compute DE against the remaining cells. scSketch compares selected cells to the non-selected background using a Welch t-test (computed from summary statistics) and displays a ranked gene list analogous to standard cluster-based DE but computed directly from interactive selections. Results are shown as an interactive table (test statistic and p-values), with gene-level plots available on click.

### 2.3 Directional sketching and statistical testing

For each user-defined direction via IDS, scSketch computes correlations between gene expression and cell projections along the sketched direction, producing a ranked list of genes associated with the trajectory (**Supplementary Figure S1**). A key challenge in exploratory analysis is multiple hypothesis testing, as each sketch involves testing thousands of genes and users may repeat this process many times [17]. To address this challenge, scSketch applies online FDR control using the LORD++ procedure to maintain statistical validity in a sequential setting [18-21]. This approach adaptively updates significance thresholds across sketches, balancing discovery and error control during iterative exploration.

### 2.4 Pathway-aware downstream analysis

To facilitate biological interpretation, scSketch integrates pathway-level context directly into the interactive workflow. For genes identified along a sketched trajectory, scSketch queries the Reactome database to retrieve pathway memberships and molecular interaction information. Pathways associated with selected genes are displayed inline, allowing users to transition seamlessly from exploratory sketches to pathway-level insight without exporting results or invoking separate enrichment tools.

*Complete algorithmic details, statistical assumptions, ordering strategies, and pseudocode for the online FDR procedure are provided in* ***Supplementary Methods S1***.

## 3 Results

### 3.1 scSketch available as a reusable software package

scSketch is provided as an open-source Python package designed to be directly imported and used within Jupyter notebook workflows. The package is distributed via PyPI, enabling straightforward installation using standard Python package managers (e.g., pip install scsketch). scSketch exposes a programmatic interface for loading single-cell datasets, launching interactive visualizations, and accessing sketch-derived results for downstream analysis. Documentation, an example notebook, and source code are available on the project’s GitHub repository (https://github.com/colabobio/scsketch).

### 3.2 Case study: differentiation of oral keratinocytes in HCMV infection

To demonstrate the utility of scSketch, we applied it to scRNA-seq data from primary HOKs infected with HCMV [11]. The dataset included cells from both mock-infected controls and cells infected with TB40eGFP [22, 23] and MOLD strains [24-26], sampled at one- and three-days post infection. Although the experiment includes discrete infection conditions, individual cells exhibited a broad and continuous range of viral transcriptional load and corresponding host responses, motivating trajectory-based analysis in addition to clustering. Using scSketch, we interactively defined trajectories across the embedding that spanned cells with low to high viral transcriptional load, enabling examination of user-guided directions of variation rather than reliance on a single precomputed trajectory. Directional correlation analysis revealed strong upregulation of interferon-stimulated genes (ISGs), including *IFI27, ISG15*, and *RSAD2*, along the infection gradient. In contrast, genes associated with mitochondrial metabolism and oxidative phosphorylation, such as *NDUFA1* and *COX6C*, were negatively correlated, suggesting a progressive viral reprogramming of host energy metabolism. Pathway-aware interpretation via Reactome further contextualized these results. Positively correlated genes were enriched for pathways related to innate immune signaling, cytokine-mediated responses, and autophagy, while negatively correlated genes mapped to oxidative phosphorylation, TCA cycle, and cell cycle regulation. These findings recapitulate known HCMV host-response strategies [11, 27-32], while illustrating how interactive, trajectory-based exploration can surface continuous biological gradients that are not readily captured by cluster-level DE alone.

### 3.3 Performance and scalability

In addition to its novel features, scSketch is designed to be efficient and scalable for modern single-cell datasets. In compute-only benchmarks (excluding interaction and rendering, which reliably present interactive framerates, and network requests), analysis runtime increased predictably with selection size and dataset sparsity. On an Apple M2 Pro laptop (16GB RAM), directional correlation testing (correlations + p-values) for a 25,531-cell selection took 0.27 s on a 161,764 cells x 20,525 genes dataset of peripheral blood mononuclear cells (PBMC161k) and 0.28 s on 1,206,761 cells x 32,357 genes dataset of cytomegalovirus infection (CMV1.2M); the subsequent online FDR update (LORD++) took 0.15 s (PBMC161k) and 0.16 s (CMV1.2M). DE for a 25,531-cell selection required a one-time global-statistics precompute of 1.92 s (PBMC161k) and ∼48.25 s (CMV1.2M), after which per-selection DE took ∼0.12 s (PBMC161k) and ∼0.13 s (CMV1.2M), increasing to 21.96 s for a 351,863-cell exported user selection in the CMV1.2M dataset. Note that runtime of directional correlation testing depends on the number of selected cells and number of genes, not total number of cells, while DE does depend on total number of cells; however, once the global statistics needed for DE are computed for a given dataset, they are cached in memory so that all subsequent DE selections do not require recalculation. Full benchmarking methodology and results are provided in **Supplementary Table S1**.

## 4. Conclusions

scSketch unifies interactive sketch-based trajectory exploration, online FDR control, and pathway-aware interpretation within a single Jupyter widget for single-cell analysis. By allowing users to define and iteratively refine hypothesis-driven trajectories directly on low-dimensional embeddings, scSketch bridges exploratory visualization with statistically rigorous inference. Applied to scRNA-seq data from HCMV-infected HOKs, scSketch recapitulated known host immune and metabolic response programs while highlighting continuous infection-associated gradients that are difficult to capture using cluster-centric analyses alone. More broadly, scSketch enables flexible interrogation of continuous structure across diverse single-cell datasets without requiring predefined trajectories or extensive parameter tuning. Future extensions will incorporate regulatory gene network analysis, support additional multimodal data types, and GPU implementation of the directional and differential analyses, further expanding scSketch’s utility as a platform for interactive, statistically principled exploration of single-cell data.

## 5. Data Availability

scSketch is open source and available as a Python package via PyPI (https://pypi.org/project/scsketch/), with source code and documentation hosted on GitHub (https://github.com/colabobio/scsketch).

## 6. Supplementary Data

Supplementary data are available at Bioinformatics online.

## 7. Acknowledgement

The authors thank Trevor Manz, the creator of anywidget that makes this work possible, as well as Thomas McCormick for useful discussions and support.

