## Supplementary Materials for "scSketch: Interactive Sketch-based Trajectory Exploration and Pathway-Aware Analysis of Single-Cell Data"

### Supplementary Methods

### S1 Online FDR Control

#### S1.1 Notation and general setup

Online hypothesis testing aims to sequentially test the (null) hypotheses $H_i : \theta_i = 0$ while maintaining a notion of online error control. For each hypothesis, we assume access to p-values that are uniformly distributed under the null; i.e., $p_i \sim Unif[0, 1]$ when $\theta_i = 0$. The hypothesis $H_i$ is rejected when $p_i \leq \alpha_i$, where the thresholds $\alpha_i$ must be determined by the online FDR controlling algorithm. We define $R_i := 1\{p_i \leq \alpha_i\}$ as the indicator that $H_i$ is rejected. A false positive at step $i$ is denoted by $V_i^\theta := 1\{p_i \leq \alpha_i \; and \; \theta_i = 0\}$. We follow the convention of [19] of appending $\theta$ to quantities that are not observed. Denote the indices of the rejected hypotheses up to $n$ at which rejections occurred by:

$$T(n) := \{i \; such \; that \; i \leq n \; and \; R_i = 1\}$$

$$:= \{\tau_1, \ldots, \tau_{R(n)}\}$$

Let $R(n) := \sum_{i=1}^{n} R_i$ and $V(n) = \sum_{i=1}^{n} V_i^{\theta_i}$ be the number of discoveries and false discoveries up to time $n$, respectively. The false discovery proportion up to time $n$ is denoted by,

$$FDP^\theta(n) := \frac{V^\theta(n)}{\max(R^\theta(n), 1)}$$

and its expectation is the false discovery rate (FDR):

$$FDR(n) := E_\theta \left[FDP^\theta(n)\right].$$

#### S1.2 Application context

The interactive workflow offered by scSketch through IDS results in two layers of hypothesis testing multiplicity. For each user-specified direction, we test a batch of G genes, and multiple directions can be tested over the course of an exploratory analysis. To frame these tests as a special case of online hypothesis testing problem, we must specify the individual hypotheses $H_i : \theta_i = 0$ and their associated p-values $p_i$.

Suppose that the embeddings are stored in the matrix $Z \in \mathbb{R}^{N\times 2}$. Suppose that after the m^th^ interaction the user has drawn the direction $v_m \in \mathbb{R}^2$ on the embedding visualization. We compute the cell-level projections $u_m = Zv_m$ and then evaluate the correlation of each gene $x_g \in \mathbb{R}^N$ with this projection: $r^m_g := Cor(x_g, u_m)$. We sequentially test the significance of these correlations. Specifically, let $i = (m - 1)G + g$ index the test that the g$^{th}$ gene is associated with the m$^{th}$ user-specified direction. The corresponding null hypothesis is $H_i : r_i = 0$. We obtain p-values for this test using a t-test with $N - 2$ degrees of freedom. Under the null and for sufficiently large $N$, these p-values are approximately uniformly distributed.

Note that, unlike standard online testing, we have control over the order in which genes are tested for any given $v_m$. To maximize power in the LORD++ algorithm, it is beneficial to have truly non-null hypotheses appear in batches. To this end, we first apply a single-linkage hierarchical clustering to the genes $[x_1, ..., x_G]$ using $1 - Cor(x_g, x'_g)$ as the distance between genes g, g′. For each m, we test genes according to the leaf order of the resulting hierarchical clustering tree.

In the current implementation of scSketch, this leaf ordering is an option in the analysis. The ordering of genes given in the data is respected, so the user can order the genes themselves using a hierarchical clustering over leaves. Future implementations may do this automatically or via a parameter provided to the scSketch object.

#### S1.3 LORD++ algorithm

Let $\gamma_i$ be a non-increasing sequence satisfying $\sum_i \gamma_i = \alpha$. The Levels based on Recent Discoveries (LORD) algorithm [19, 20] sets thresholds according to,

$$\alpha_n = w_0\gamma_n + b_0\left(\sum_{l \in T(n)} \gamma_{n-l}\right)$$

where $w_0$ and $b_0$ are algorithm hyperparameters, and $l$ the index of the last discovery. For appropriate $w_0$, $b_0$, this algorithm can be shown to control $FDR^{\theta}(n)$ at all $n$ under appropriate assumptions on the dependence structure between $p_i$. The first term on the right can be interpreted as a series of decreasing thresholds that makes rejection more difficult over time. The second term can be interpreted as a relaxation of this threshold when many discoveries have been made, especially if those discoveries have been recent. Indeed, the more discoveries that are made, the larger the index set $T(n)$ of the sum. This increases $\alpha_n$, which makes rejection easier. Further, when discoveries are recent, the indices $n - l$ in the summation will be smaller, and since $\gamma_n$ is non-increasing the associated $\gamma_{n-l}$ will be larger. Hence, the more recent discoveries increase the threshold $\alpha_n$ more than old discoveries.

scSketch adopts a variation on LORD called LORD++ [21]. This algorithm observes that, by distinguishing between the first rejection from all subsequent rejections, the power of LORD can be strictly improved. Formally, the decision rule divides into two cases, depending on whether there has only been one discovery up to time n. In the case that only one discovery has been made so far, the rule is

$$\alpha_n = w_0\gamma_n + (\alpha - w_0)\gamma_{n-\tau_1}$$

but if two or more discoveries have already been made by time n, then the rule becomes,

$$\alpha_n = w_0\gamma_n + (\alpha - w_0)\gamma_{n-\tau_1} + \left(\sum_{l \in T(n)-\{\tau_1\}} \gamma_{n-l}\right)\alpha.$$

Note that these thresholds have the same qualitative form as the original LORD thresholds, except that $\gamma_{n-\tau_1}$ has been isolated from the rest of the sum and that the free parameter $b_0$ has been removed. Like the original LORD procedure, this series of thresholds leads to online FDR control under appropriate conditions on $w_0$ and p-value dependence.

Following the implementation [15, 16] by the original algorithm developers, we set default hyperparameters:

$$w_0 = 0.005$$

$$\gamma_n = 0.0722\frac{log\ (n \vee 2)}{n \cdot exp\ \left(\sqrt{log\ (n)}\right)}$$

### S2 Directional Analysis Algorithm (implementation)

scSketch's directional analysis takes two inputs: (i) a gene expression matrix X (cells × genes) and (ii) a two-dimensional embedding E (cells × 2; e.g., UMAP). For a single user sketch, scSketch defines a set of selected cells $C_1,\ldots,C_n$ and restricts all computations to those cells.

**Directional projection.** Let v denote the user-specified direction in the embedding (**Figure S1**). scSketch assigns each selected cell $C_i$ a scalar projection $p_i$ by projecting its embedding coordinate onto v (i.e., $p_1, \dots, p_n$ in **Figure S1**).

**Gene–direction association.** For each gene g, let $(G_{g1}, \dots, G_{gn})$ denote that gene's expression values across the selected cells. scSketch computes a Pearson correlation between the projection vector and the gene's expression vector:

$$R_g = pearson((p_1, \dots, p_n), (G_{g1}, \dots, G_{gn})),$$

which is the basis for gene ranking (**Figure S1**). Two-sided p-values are computed using the standard correlation t-test with $df = n - 2$.

To support sparse scRNA-seq matrices without densifying X, scSketch computes correlations from summary statistics on the selected cells (gene-wise sums, sums-of-squares, and X^T multiplied by the centered projection vector). If n < 3, the projections have near-zero variance, or a gene has zero variance within the selection, scSketch returns $R_g = 0$ and a non-significant p-value for numerical stability. The resulting p-values are then passed to the online FDR procedure described in **Supplementary Methods S1**.

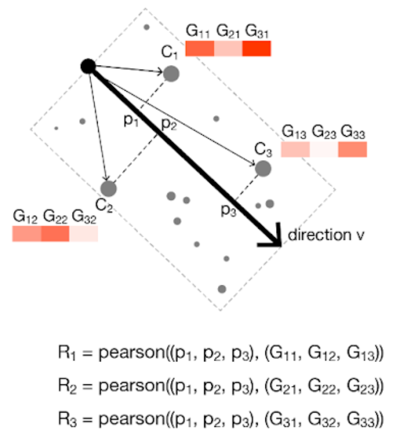


**Supplementary Figure S1.** Directional analysis in scSketch. Cells selected by the user ($C_1, \dots, C_n$) are projected onto the sketched direction v to obtain a one-dimensional coordinate ($p_1, \dots, p_n$). For each gene g, scSketch computes $R_g = pearson((p_1, \dots, p_n), (G_{g1}, \dots, G_{gn}))$ and ranks genes by their association with the direction.

Alt text: Schematic of scSketch directional analysis. A subset of cells selected on a 2D embedding ($C_1, \dots, C_n$) is projected onto a user-drawn direction vector v to produce a list of their respective 1D coordinates ($p_1, \dots, p_n$). For each gene g, scSketch computes the Pearson correlation between gene expression across the selected cells ($G_{g1}, \dots, G_{gn}$) and the projection values, yielding $R_g$, and ranks genes by strength of association with the sketched direction.

### S3 Performance Test (compute-only)

We tested scSketch runtime for the two core computational modules - Directional Search (DS) and Differential Expression (DE) - on an Apple M2 Pro laptop (10-core CPU, 16GB RAM). Benchmarks were run as compute-only tests (excluding IDS' visualization/rendering, gene-click plotting, and network queries to external databases). For DS, we timed (i) correlation plus p-value computation across all genes and (ii) the online FDR update step (LORD) separately. For DE, we timed (i) a one-time per-dataset global precomputation of gene-wise sums and sums-of-squares across all cells (to enable fast repeated DE queries) and (ii) per-selection Welch t-tests computed from summary statistics for selected vs. background cells. The full results from the compute-only tests are provided in the table below.

### Supplementary Table S1

***Supplementary Table S1.*** *Compute-only runtime benchmarks for scSketch's Directional Search (DS) and Differential Expression (DE) modules across three dataset scales (PBMC3k, PBMC161k, and CMV1.2M) on an Apple M2 Pro laptop (10-core CPU, 16GB RAM). For each dataset and selection size (n_selected), the table reports dataset dimensions (n_cells, n_genes), matrix sparsity (X_nnz_total, X_sel_nnz), and runtimes in seconds for DS correlation + p-value computation (dir_corr_p_s), the LORD++ online FDR update (dir_lord_s), the one-time DE global summary-statistics precomputation (de_global_stats_s), and per-selection DE computation (de_compute_s). LORD++ is computed with a Numba-accelerated implementation when available (with a pure-Python fallback), which can introduce a one-time JIT compilation overhead on the first call in a fresh Python process; subsequent LORD++ updates run in compiled code.*

| dataset | n_cells | n_genes | X_nnz_total | n_selected | X_sel_nnz | dir_corr_p_s | dir_lord_s | de_global_stats_s | de_compute_s |
|---|---|---|---|---|---|---|---|---|---|
| pbmc3k | 2638 | 1838 | | 500 | | 0.0033 | 0.0409 | 0.0123 | 0.0101 |
| pbmc3k | 2638 | 1838 | | 1000 | | 0.0057 | 0.0434 | 0.0123 | 0.0060 |
| pbmc3k | 2638 | 1838 | | 2000 | | 0.0112 | 0.0919 | 0.0123 | 0.0113 |
| pbmc161k | 161764 | 20525 | 336245756 | 20661 | 46461966 | 0.2839 | 0.1583 | 1.9170 | 0.1132 |
| pbmc161k | 161764 | 20525 | 336245756 | 25531 | 57546958 | 0.2701 | 0.1522 | 1.9170 | 0.1240 |
| pbmc161k | 161764 | 20525 | 336245756 | 50000 | 113143093 | 0.6867 | 0.1621 | 1.9170 | 0.5579 |
| cmv1.2m | 1206761 | 32357 | 2056839573 | 25531 | 37917401 | 0.2777 | 0.1569 | 48.2472 | 0.1263 |
| cmv1.2m | 1206761 | 32357 | 2056839573 | 143672 | 214397116 | 1.6847 | 0.2008 | 48.2472 | 1.6984 |
| cmv1.2m | 1206761 | 32357 | 2056839573 | 376364 | 572020646 | 10.7304 | 0.2350 | 48.2472 | 14.1228 |

In summary, across three dataset scales - PBMC3k (2,638 cells x 1,838 genes), PBMC161k (161,764 x 20,525; WNN UMAP), and the CMV1.2M dataset (1,206,761 x 32,357; UMAP) - DS correlation plus p-value computation scaled with selection size: on PBMC161k it ranged from 0.27-0.69 s for selections of 20,661-50,000 cells, and on the CMV1.2M dataset it ranged from 0.28 s (25,531 cells) to 10.73 s (376,364 cells). The LORD update step took consistently longer than the correlation step and depended primarily on the number of genes tested, ranging from 0.15-0.16 s on PBMC161k and 0.16-0.24 s on the CMV1.2M dataset. DE showed two regimes: a one-time global-statistics precomputation cost of 1.92 s (PBMC161k) and 48.25 s (CMV1.2M), followed by per-selection DE times that scaled with selection size (PBMC161k: 0.11-0.56 s for 20,661-50,000 selected cells; CMV1.2M: 0.13-14.12 s for 25,531-376,364 selected cells). To complement standardized synthetic selections, we also benchmarked a real user-defined selection exported from a scSketch session on the CMV1.2M dataset (351,863 selected cells): DS correlation plus p-values took 11.24 s, LORD update took 0.35 s, and per-selection DE took 21.96 s (with one-time DE precomputation 53.80 s). Together, these results characterize scSketch's compute-time scaling for interactive trajectory probing and DE on datasets up to ~1.2M cells under a reproducible benchmark protocol. Thanks to the Numba optimizations, most of the calculations result in very short wait times for the directional and differential gene rankings, minimizing disruption of the interactive workflow; only in very large datasets (several millions of cells) the runtime of these calculations result in noticeable delays, but even then, they do not affect the IDS process itself as they are performed only upon user request after directional sketching is completed. We will explore GPU implementations of all gene ranking calculations as well as techniques such as subsampling in future versions of scSketch to reduce waiting times and enable seamless interactive analyses even for extremely large single-cell datasets.
